# A high-throughput COPD bronchosphere model for disease-relevant phenotypic compound screening

**DOI:** 10.1101/2022.12.16.520302

**Authors:** Pranjali Beri, Young Jae Woo, Katie Schierenbeck, Kaisheng Chen, S. Whitney Barnes, Olivia Ross, Douglas Krutil, Doug Quackenbush, Bin Fang, John Walker, William Barnes, Erin Toyama

## Abstract

COPD is the third leading cause of death worldwide, but current therapies for COPD are only effective at treating the symptoms of the disease rather than targeting the underlying pathways that are driving the pathogenic changes. The lack of targeted therapies for COPD is in part due to a lack of knowledge about drivers of disease progression and the difficulty in building relevant and high throughput models that can recapitulate the phenotypic and transcriptomic changes associated with pathogenesis of COPD. To identify these drivers, we have developed a cigarette smoke extract (CSE)-treated bronchosphere assay in 384-well plate format that exhibits CSE-induced decreases in size and increase in luminal secretion of MUC5AC. Transcriptomic changes in CSE-treated bronchospheres resemble changes that occur in human smokers both with and without COPD compared to healthy groups, indicating that this model can capture human smoking signature. To identify new targets, we ran a small molecule compound deck screening with diversity in target mechanisms of action and identified hit compounds that attenuated CSE induced changes, either decreasing spheroid size or increasing secreted mucus. This work provides insight into the utility of this bronchosphere model in examining human respiratory diseases, the pathways implicated by CSE, and compounds with known mechanisms of action for therapeutic development.

## Introduction

Chronic obstructive pulmonary disease (COPD) is the third leading cause of death worldwide, accounting for approximately 6% of global deaths in 2019(1). It is characterized by irreversible airflow obstruction and persistent inflammation, most commonly in response to cigarette smoke exposure(2–5). The airways of COPD patients are characterized by structural changes to the airway epithelial cells that contribute to airflow obstruction, such as: increased mucus production and secretion (particularly MUC5AC), decreased mucus clearance through ciliated cell dysfunction, and decreased ion channel activity leading to decreased airway hydration which normally thins mucus and makes it easier to clear(3,6,7). The mechanisms behind these phenotypic changes are still not well understood, and current treatments of COPD mainly consist of corticosteroids and bronchodilators that aim to relax the airway to improve airflow(8). To enable the discovery of novel and effective therapies for COPD, we need scalable high-throughput disease models that can capture the relevant complex phenotypic and transcriptomic changes associated with the disease.

Since the primary cause of COPD in patients is cigarette smoking, a commonly utilized method of inducing a COPD-like phenotype *in vitro* is to expose airway epithelial cells to cigarette smoke in one of three formats: cigarette smoke condensate (CSC)(9), cigarette smoke extract (CSE)(10), or whole smoke (WS)(11). Often these models are not well suited for therapeutic drug discovery efforts for three main reasons. One, they lack sufficient complexity because they do not utilize differentiated bronchial epithelial cells, which is necessary to recapitulate the mixture of cellular subpopulations that make up the airway epithelium (12–14). Two, they do not investigate the effects of long-term exposure to cigarettes, often treating for shorter bursts which models acute exposure but does not reveal longer term changes such as mucus secretion shifts(14,15). Or three, the more complex models are often constructed within air-liquid interface plates. These models are not scalable for use in high-throughput screening which requires at least a 384-well plate format to enable the screening of larger small molecule libraries that are typical of a drug discovery pipeline (10,11,16,17). Airway organoids, or bronchospheres, are 3D self-assembling structures that can be differentiated from primary human airway basal cells, where the mature bronchospheres are composed of distinct subpopulations of cells present in the airways epithelium, such as ciliated cells and secretory cells (i.e. goblet cells)(18). They are advantageous in that they can be scaled down to 384-well plate formats and do not require air-liquid interface plates^16,17^. However, a COPD-mimetic disease model using these bronchospheres has not been previously generated. While bronchospheres could be generated from COPD patient-derived basal cells, it is advantageous to induce a disease phenotype in healthy cells to understand key disease drivers while minimizing donor-to-donor variability introduced by the inherent heterogeneity of COPD(19,20).

Here we present a 384-well *in vitro* bronchosphere model that recapitulates key aspects of the COPD disease phenotype. Treatment of human bronchial epithelial cell (HBEC)-derived bronchospheres to 3% CSE over one, two, and three weeks of treatment resulted in a decrease in bronchosphere size, which is an indicator of loss of swelling or hydration state in the lumen(21–23), and an increase in luminal secretion of MUC5AC across all timepoints, providing two separate phenotypic readouts of responses to CSE. Bulk RNAseq analysis shows an overlap in the differentially expressed genes (DEGs) in the *in vitro* model with comparisons from three independent patient smoker vs non-smoker datasets, and DEGs in the CSE-treated bronchospheres show enrichment in COPD-relevant GO terms, KEGG pathways, and Human MSigDB signatures. We used this reproducible, disease-relevant model in a diverse mechanism of action (MoA) compound screen where hit compounds were able to attenuate the two key phenotypic changes. Overall, this study has yielded insights into the pathways that are involved in the aberrant phenotype that presents after cigarette smoke exposure and can lead to potential therapeutic avenues in the treatment of COPD.

## Results

### Bronchospheres treated with CSE exhibit reduced spheroid area and increased MUC5AC secretion across all timepoints

To generate a high-throughput disease model for COPD drug discovery, bronchospheres derived from HBECs from three independent healthy donors were first cultured in 384-well plates in 2.5D culture on Matrigel. We found that co-culture with an NIH 3T3 feeder cell population in the plate bottom resulted in larger spheroids with more obvious lumen formation (Figure 1A and Supplemental Figure 1A). Previous studies have shown that differentiation of bronchosphere cultures for two weeks is sufficient to generate distinct cellular subtypes(18,24) (Supplemental Figure 1B), so bronchosphere cultures were differentiated for two weeks prior to continuous CSE treatment for 1, 2, or 3 weeks (Figure 1B) with either 3% CSE or 0% CSE control. At each time point, bronchosphere cultures were imaged live to observe changes in size due to CSE treatment, then fixed and stained for expression of secreted mucins (MUC5AC and MUC5B) in the lumen (Figure 1B).

**Figure 1:**
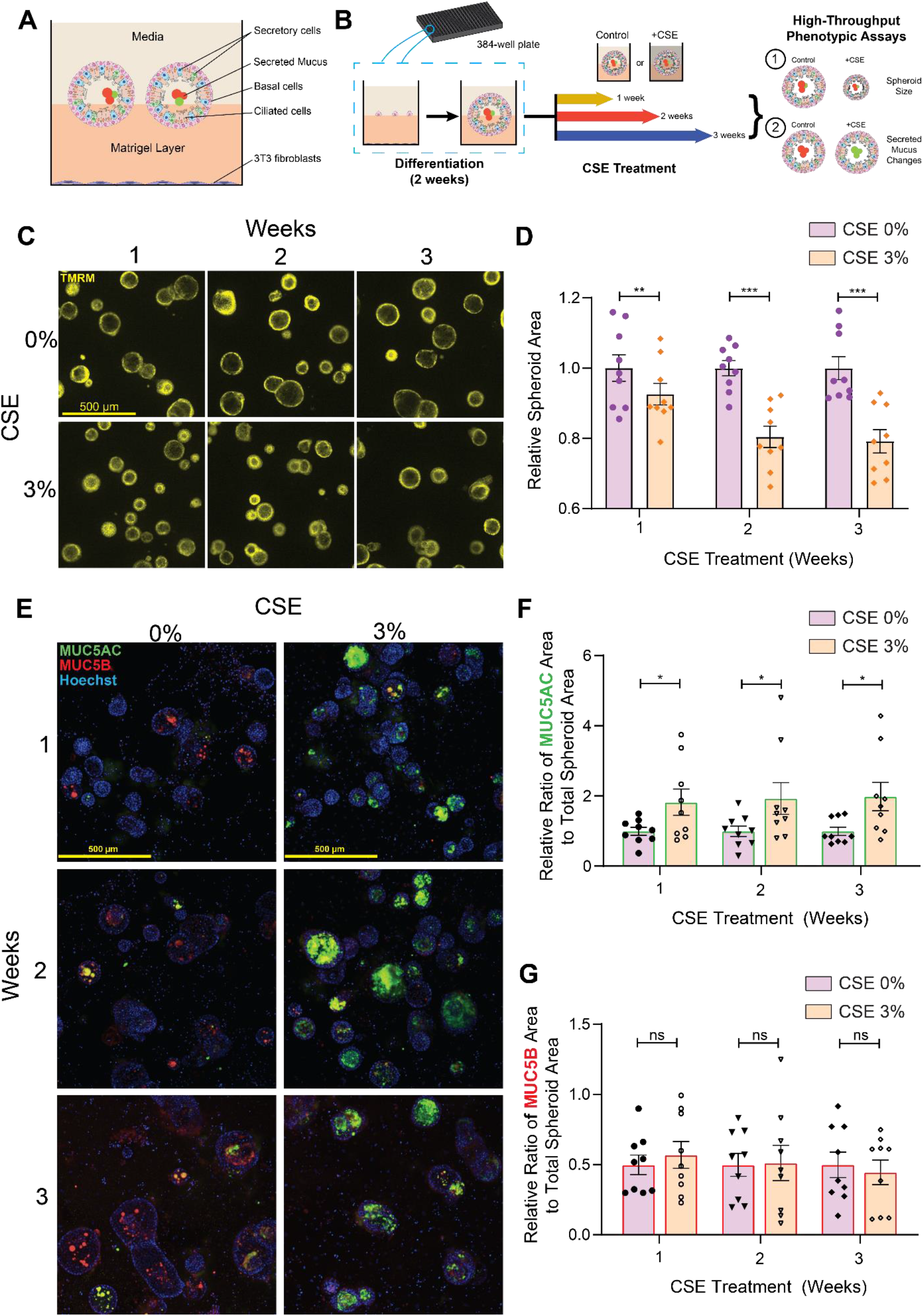
CSE-treated bronchospheres have decreased spheroid area and increased MUC5AC secretion in the lumen compared to control. **A)** Schematic of a well within a 384-well plate with bronchosphere organoids in 2.5D culture with a 3T3 feeder cell layer at the well bottom. **B)** Schematic of the experimental workflow. Bronchospheres were seeded and differentiated over two weeks, after which they were exposed to 0 or 3% CSE for 1-, 2-, and 3-week continuous treatment. At each timepoint, spheroids were imaged live to assess changes in size, then fixed and stained for secreted mucins. **C)** Representative images of live bronchospheres stained with TMRM dye that show changes in area with CSE treatment. **D)** Bar plot of spheroid area at all timepoints treated with 0 or 3% CSE. Conditions are comprised of 3 biological replicates from 3 independent donors. Datapoints were normalized to the average of the control condition for each donor and at each timepoint, due to donor-to-donor variability. Data is represented as mean ± SEM and each timepoint was analyzed by paired two-tailed t-test, **p<0.01, ***p<0.001. **E)** Representative images of bronchospheres stained for MUC5AC and MUC5B that show increased MUC5AC secretion in the bronchospheres lumen with CSE treatment. **F)** Bar plot of ratio of MUC5AC and **G)** MUC5B cluster area to total spheroid area. Conditions are comprised of 3 biological replicates from 3 independent donors. Datapoints were normalized to the average of the control condition for each donor and at each timepoint, due to donor-to-donor variability. Data is represented as mean ± SEM and each timepoint was analyzed by paired two-tailed t-test, *p<0.05.

CSE treatment resulted in a significant decrease in bronchosphere size at all time points, with greater decreases occurring after 2 and 3 weeks of treatment (Figure 1C and D). Since an important disease phenotype of COPD is the increased secretion of airway mucus, particularly MUC5AC(10,24), we wanted to demonstrate that this change can be captured in the CSE-treated bronchospheres. To quantify changes in MUC5AC, we calculated the change in area of the secreted mucin clusters within the bronchosphere lumen and divided by the total spheroid area, thus obtaining the ratio of the total spheroid area occupied by the mucin cluster. We found that bronchospheres treated with CSE had an increase in luminal MUC5AC ratio compared to control at all three time points (Figure 1E-F), while there was no change in the MUC5B ratio relative to control (Figure 1G). These data indicate that the CSE-treated bronchospheres undergo multiple phenotypic changes in response to CSE-treatment that can be quantified and utilized in phenotypic screening applications.

### Bronchosphere model treated with CSE captures smoking signature in humans

To further investigate how well the model recapitulates the airways of advanced smokers, we isolated RNA from bronchospheres treated with 0 or 3% CSE for 1, 2, or 3 weeks and performed bulk RNA sequencing and analysis to calculate CSE treatment dependent differentially expressed genes (DEGs) (Figure 2A). In addition to time-point specific DEGs, DEGs after pooling all time points were calculated to maximize power in identifying the CSE-treated bronchosphere signature using mixed model (See Methods for more detail). With FDR controlled at 5% and DEGs further restricted to a greater than 2 fold-change difference, there were 408 DEGs in the pooled analysis, 437 DEGs in 1 week treated alone, 446 DEGs in 2 weeks treated alone, and 510 DEGs in 3 weeks treated alone (Figure 2A and Supplemental Table S2).

**Figure 2:**
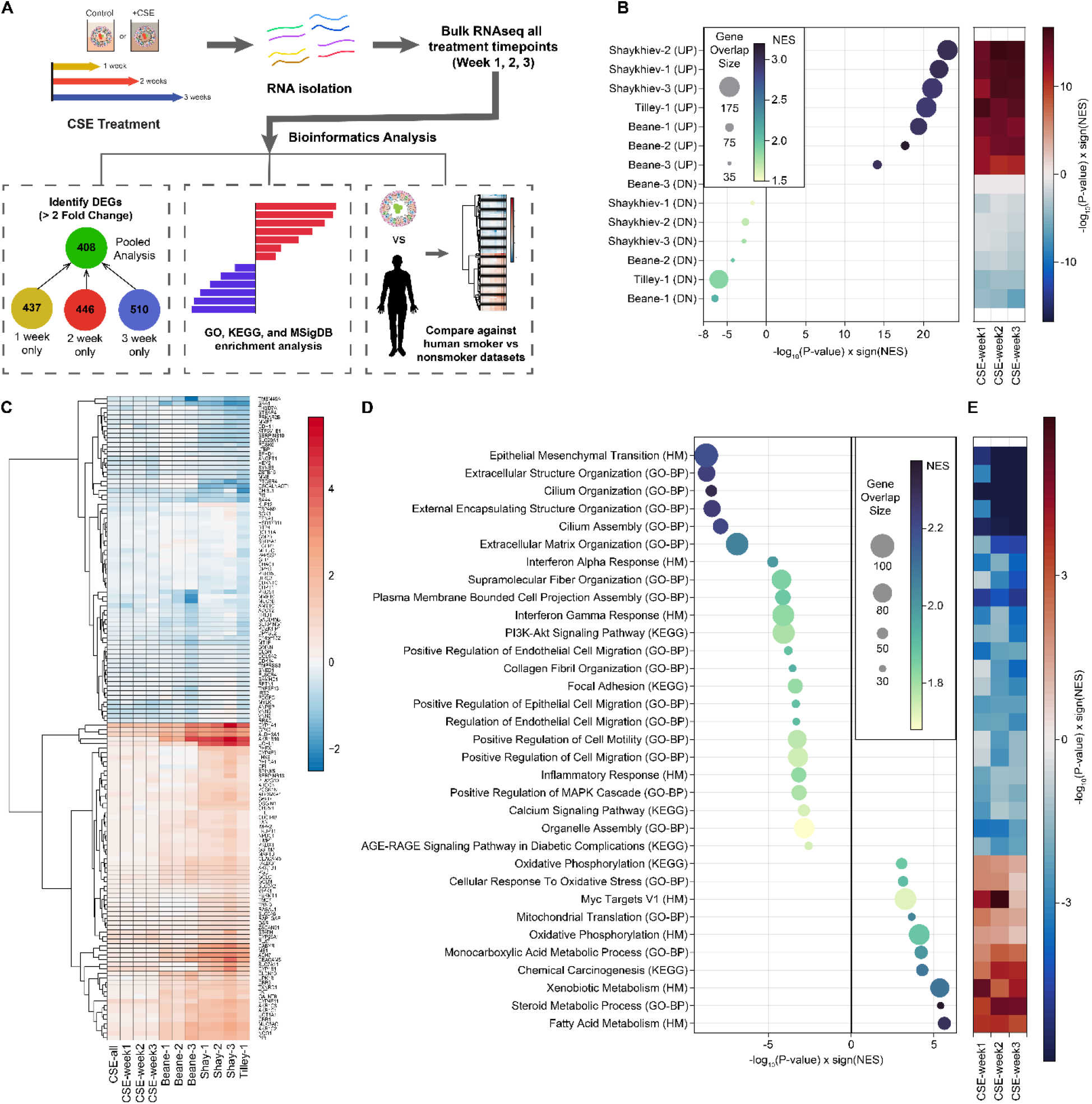
CSE-treated bronchospheres capture smoke signatures in humans. **A)** Schematic of RNA sequencing and bioinformatics analysis. The number of DEGs at each timepoint and pooled timepoints were calculated with FDR controlled at 5% and fold-change >2. **B)** Enrichment of up- and down-regulated genes from CSE-treated bronchospheres in smoking signatures comprised of 7 different comparisons generated from 3 separate datasets (a legend of the comparisons is provided in Supplemental Table S6). The heatmap on the right indicates enrichment at each timepoint with the 7 dataset comparisons. As expected, a positive enrichment score with the UP comparisons indicates that upregulated genes in the CSE-bronchospheres are enriched in the upregulated genes of the human signatures, while a negative enrichment score with the DN comparisons indicates that the downregulated genes in the CSE-treated bronchospheres are enriched in the downregulated genes of the human signatures. **C)** Heat map visualization of gene expression changes (β estimates) of 75 top up-regulated and 75 top down-regulated genes from Tilley-1 comparison against the other patient comparisons as well as in the in vitro datasets, both pooled and individual timepoints. **D)** GSEA shows up- and down-regulated pathways in CSE-treated bronchospheres. Significantly associated pathways with gene-set overlap size larger than 30 genes were selected for visualization.

Smoking signatures from human samples were generated from three published datasets (See Methods)(25–27). For GSE20257 by Shaykhiev et al.(25), three comparisons were tested: 1) excluding COPD, smokers were compared against non-smokers (Shaykhiev-1), 2) including COPD, smokers were compared against non-smokers (Shaykhiev-2), and 3) Non-smokers compared against COPD smokers (Shaykhiev-3). For GSE7895 by Beane et al.(26), three comparisons were examined: 1) linear effect of smoking dependencies in the order of never smokers, former smokers, and current smokers (Beane-1), 2) never smokers vs current smokers (Beane-2), and 3) former smokers vs. current smokers (Beane-3). For GSE63127 by Tilley et al.(27), smokers were compared against non-smokers (Tilley-1). Up and down-regulated DEGs were calculated for each comparison (Supplemental Table S3) and Gene-Set Enrichment Analysis (GSEA) was performed with the hypothesis that up-regulated DEGs from CSE-treated bronchosphere models will be enriched in the up-regulated genes from human smoking data, and vice versa (Supplemental Table S4). As expected, up-regulated genes in all seven smoking signatures were significantly associated with positive enrichment scores, suggesting that DEGs up-regulated in the bronchospheres upon CSE treatment is over-represented in genes increased in the smoking cohorts (Figure 2B). To a smaller degree, DEGs down-regulated in the bronchospheres after CSE treatment were also enriched in the down-regulated smoking signatures (Figure 2B). Seventy-five top up-regulated and seventy-five top down-regulated genes from Tilley-1 comparison were selected to visualize gene expression patterns across the other six smoking comparisons using human data and four CSE-treatment comparisons using bronchosphere model (Figure 2C). The Tilley dataset was prioritized in gene list selection because it had the largest sample size; therefore, its biology is most likely generalizable amongst all other smoking effect analyses. While the effect sizes from the human smoking signature were larger than the CSE treated bronchosphere model, the direction of effects across all datasets were consistent (Figure 2C). CYP1A1(28), GPX2(29), ALDH3A1(30), and CYP4F11(31) were the top up-regulated genes and were strongly associated with responses to CSE, which further bolstered that the bronchosphere model with CSE treatment is a good *in vitro* representation of human smoking trait. Among down-regulated genes, GLIS3 was a known COPD risk gene from GWAS(32,33), and MMP7 has a known promoter polymorphism associated with early onset COPD(34). Of note, we also observed that CFTR was downregulated in the CSE-treated bronchospheres, which can, in part, explain the observed loss of size with CSE treatment(6,21,22) (Supplemental Table S2).

GSEA was performed to understand which biological pathways were significantly impacted by CSE treatment in the bronchosphere model (Supplemental Table S5). Epithelial Mesenchymal Transition (EMT) from MsigDB Hallmark (P=1.6×10^-9^), Extracellular Structure Organization (P=1.7×10^-9^) and Extracellular Matrix (ECM) Organization (P=1.2×10^-7^) were enriched with down-regulated genes (Figure 2D-E). The enrichment of down-regulated genes in these pathways increased with weeks after CSE treatment (Figure 2D-E). Cilium Organization (P=3.4×10^-9^), Cilium Assembly (P=1.2×10^-8^), Supramolecular Fiber Organization (P=5.9×10^-5^), Positive Regulation of Epithelial Cell Migration (P=4.4×10^-4^), Positive Regulation of Cell Motility (P=5.3×10^-4^) were also enriched in down-regulated DEGs, suggesting reduced cell mobility upon CSE treatment (Figure 2D-E). IFN-*α* Response (P=1.8×10^-5^), IFN-*γ* Response (P=7.7×10^-5^), and Inflammation Response (P=6.6×10^-4^) were also enriched with down-regulated DEGs (Figure 2D-E). This suggests that ECM reorganization(35), cell mobility(36), and IFN responses(37) may be negatively impacted by CSE treatment. On the other hand, up-regulated DEGs were overrepresented in Fatty Acid Metabolism (P=2.3×10^-6^), Steroid Metabolic Process (P=3.9×10^-6^), Xenobiotic Metabolism (P=4.4×10^-6^), suggesting increased metabolism upon CSE treatment (Figure 2D-E). From ClinVar 2019, primary ciliary dyskinesia was the only disease associated with DEGs in the bronchosphere model after 5% FDR correction (P=3.7×10^-8^, NES=-2.2).

### Phenotypic screen with a small molecule diversity deck identifies compounds that attenuate the size and mucus changes induced by CSE treatment

To demonstrate the phenotypic screening potential of this system and identify pathways that are involved in the decreased spheroid size and increased luminal mucus due to CSE treatment, we ran a limited small molecule screen on bronchospheres treated with CSE for two weeks (Figure 3A and Supplemental Table S7). Compounds used in the screen make up a diversity deck consisting of 301 non-proprietary compounds selected to hit a broad range of targets to identify pathways that might be responsible for the observed phenotypic changes. Compound treatment began at day 7 of CSE treatment, where bronchospheres were exposed to either DMSO or compounds at 1 or 10μM final concentrations for the remaining 7 days of CSE treatment. Bronchospheres were imaged live at the 2-week endpoint then fixed and stained for MUC5AC ratio changes in the bronchosphere lumen. After applying filters to remove compounds that cause toxicity or worsen the phenotype in the other screen, 26 final hits were identified that either decreased MUC5AC ratio (Figure 3B and Supplemental Table S8) or increased spheroid size (Figure 3C and Supplemental Table S8).

**Figure 3:**
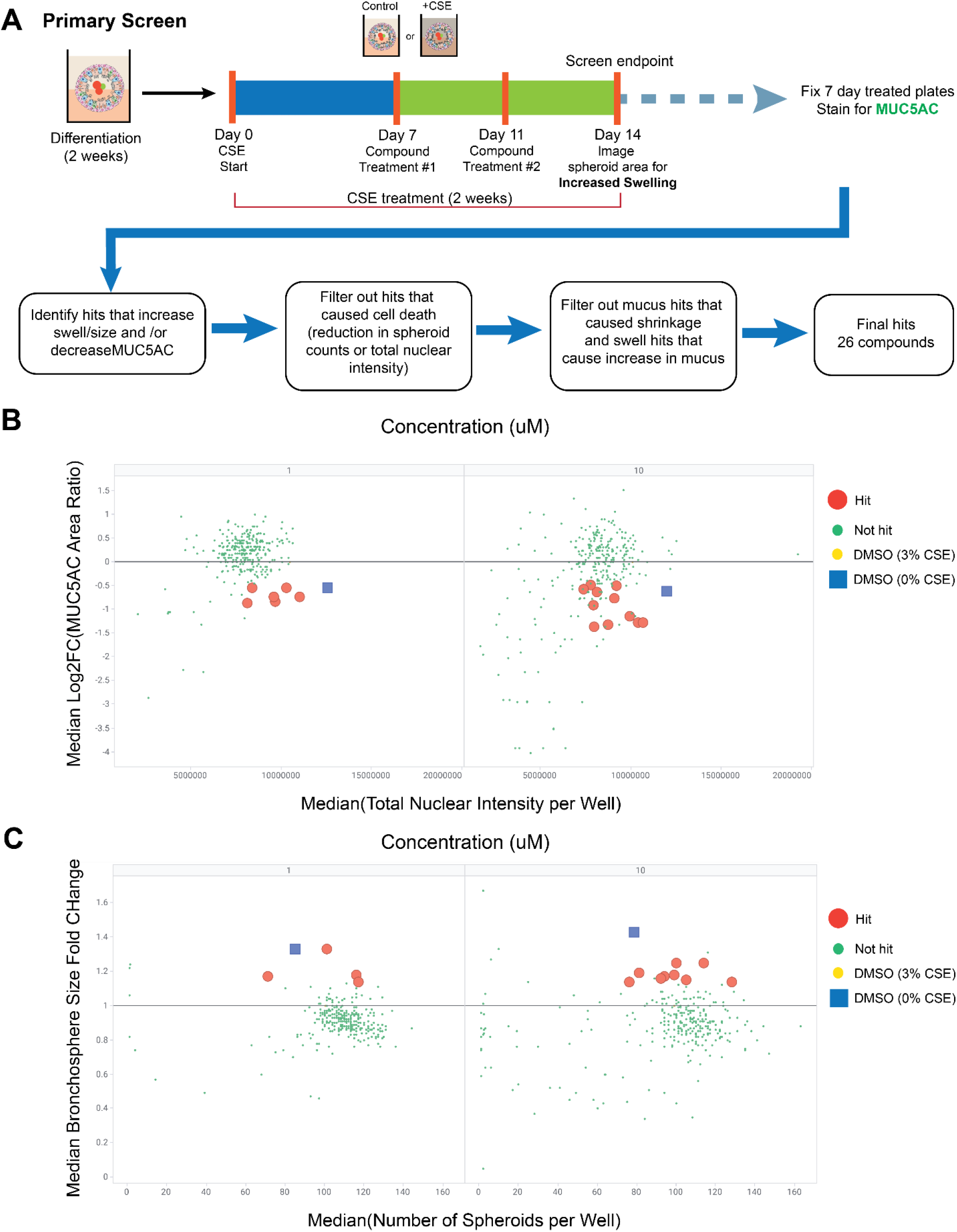
Primary small molecule screen to identify attenuators of CSE-induced decrease in size and increase in MUC5AC. **A)** Schematic of primary screen set-up, endpoints, and identification of hits. Compounds were tested at 1 and 10μM concentrations with n=3 replicates per compound and concentration. The deck consisted of 301 compounds of diverse mechanisms of action. Hit compounds at 1 and 10μM concentrations that **B)** increased spheroid size (swell) or **C)** decreased MUC5AC staining in the lumen relative to DMSO and CSE 3%-treated bronchospheres. A detailed explanation of hit selection for both readouts is provided in the methods.

To validate the primary hits, a second screen was run with the same experimental parameters to identify which compounds recapitulated their effects. Of all the compounds that restored spheroid size in the primary screen, only Compound 206 (1μM, HDAC4/5 inhibitor) validated in the secondary screen (Figure 4A-B). Meanwhile, of the compounds that reduced MUC5AC in the primary screen, Compounds 186 (1μM, SMN2 splicing modulator), 16 (10μM, NPY5R inhibitor), 41(10μM, EGFR inhibitor), 158(10μM, PIKfyve inhibitor), 243(10μM, CBP/EP300 inhibitor), and 253(10μM, HDAC6 inhibitor) validated in the secondary screen (Figure 4C-D). These hits were identified by filtering out the compounds that caused toxicity as monitored by a significant decrease in number of spheroids (Supplemental Figure 3A-B) and/or a significant decrease in the stained total nuclear intensity (Supplemental Figure 3C-D). Compounds that worsened the phenotype in the other screen were also filtered out (Supplemental Figure 4).

**Figure 4:**
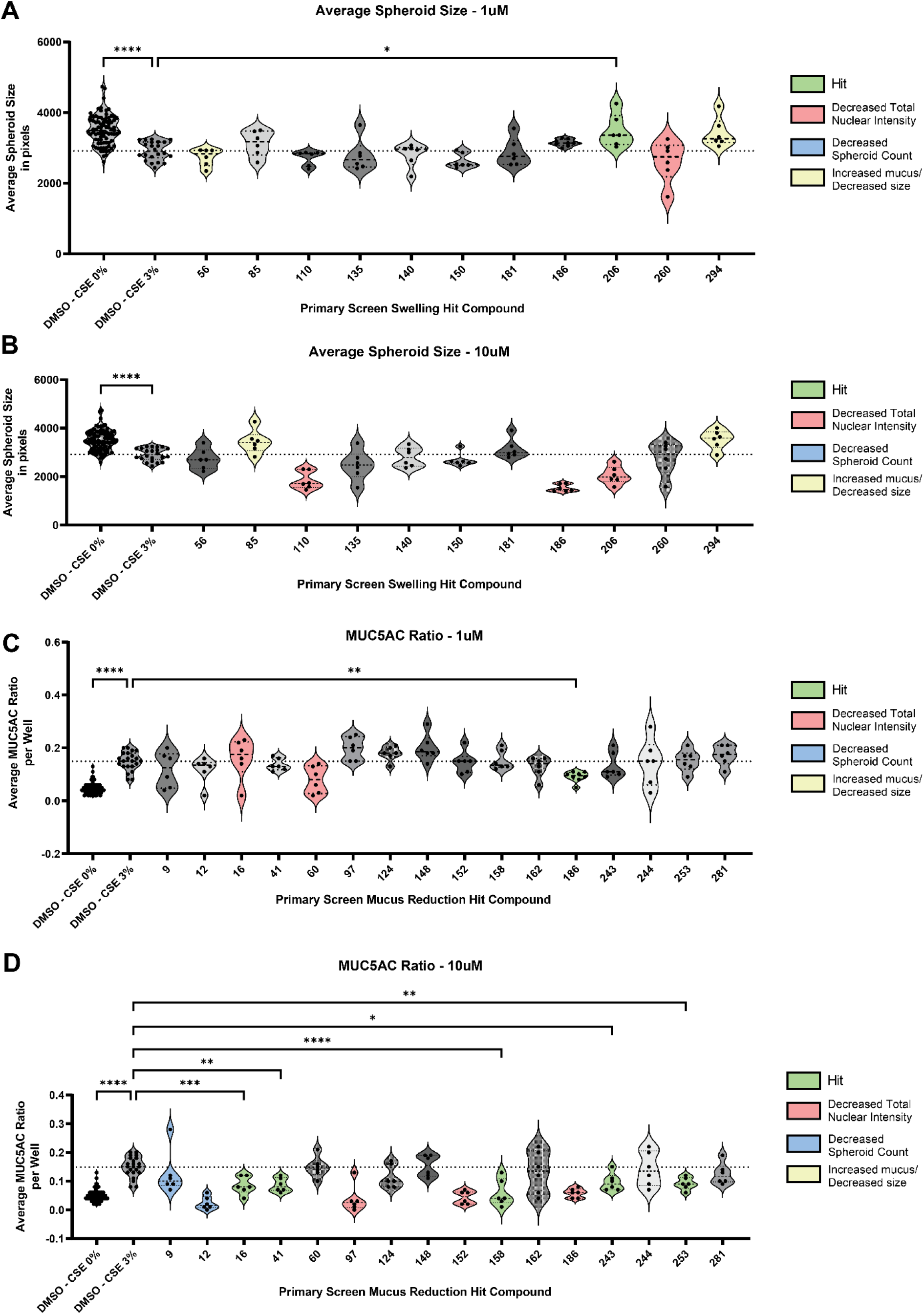
Validation of primary screen hits reveals compounds that recapitulate increased spheroid size (swell) or decreased MUC5AC in the lumen. Compounds that were identified in the primary screen as modulators of spheroid size/swell were tested at **A)** 1μM and **B)** 10μM concentrations to see if they would recapitulate the phenotype. Compound 206 (1μM) significantly increased spheroid size relative to DMSO + CSE 3% control bronchospheres. Compounds that were identified in the primary screen as modulators of MUC5AC reduction were tested at **C)** 1μM and **D)** 10μM concentrations to see if they would recapitulate the phenotype. Compounds 186 (1μM), 16 (10μM), 41(10μM), 158(10μM), 243(10μM), and 253(10μM) significantly reduced MUC5AC ration within the spheroid lumen compared to DMSO + CSE 3% control bronchospheres. Compounds that appeared to cause toxicity were identified by observing a significant decrease in spheroid counts in the swell readout or a significant decrease in the total nuclear intensity of the MUC5AC ratio readout. These data are reported in Supplemental Figure 3. Compounds that correctly attenuated the phenotype but produce the opposite phenotype in the other readout (i.e., increased spheroid area but also significantly increased MUC5AC ratio in the lumen) were also filtered out. These data are reported in Supplemental Figure 4. All individual data points represent biological replicates. All plots were analyzed by ordinary one-way ANOVA with Dunnett’s multiple comparisons test. *p<0.05; **p<0.01, **p<0.01, ***p<0.001, ****p<0.0001.

To ensure that the identified hits were not causing unforeseen phenotypic changes to the bronchospheres that were not identified in the quantitative filtration process, TMRM live dye-stained bronchosphere images were compiled and compounds that induced aberrant phenotypes compared to DMSO + 3% CSE control bronchospheres were manually filtered (Supplemental Figure 5). An unhealthy phenotype was one where there was an obvious loss of central lumen, uncharacteristic clumping, or rough spheroid boundaries. The final list of hits and their restoration of swell or mucus secretion is shown in Figure 5. An inhibitor of HDAC4 and HDAC5 (LMK235) was found to attenuate the decrease in spheroid size due to CSE treatment, while inhibitors of CBP/EP300 (CPI637), EGFR (AEE788), and HDAC6 (CHEMBL3415627) were found to attenuate the increase in MUC5AC in the bronchosphere lumen due to CSE treatment (Main Table 1). Taken together, these data show that this platform can be utilized to identify key pathways that are involved in disease-associated changes caused by exposure to byproducts of cigarette smoke.

**Figure 5:**
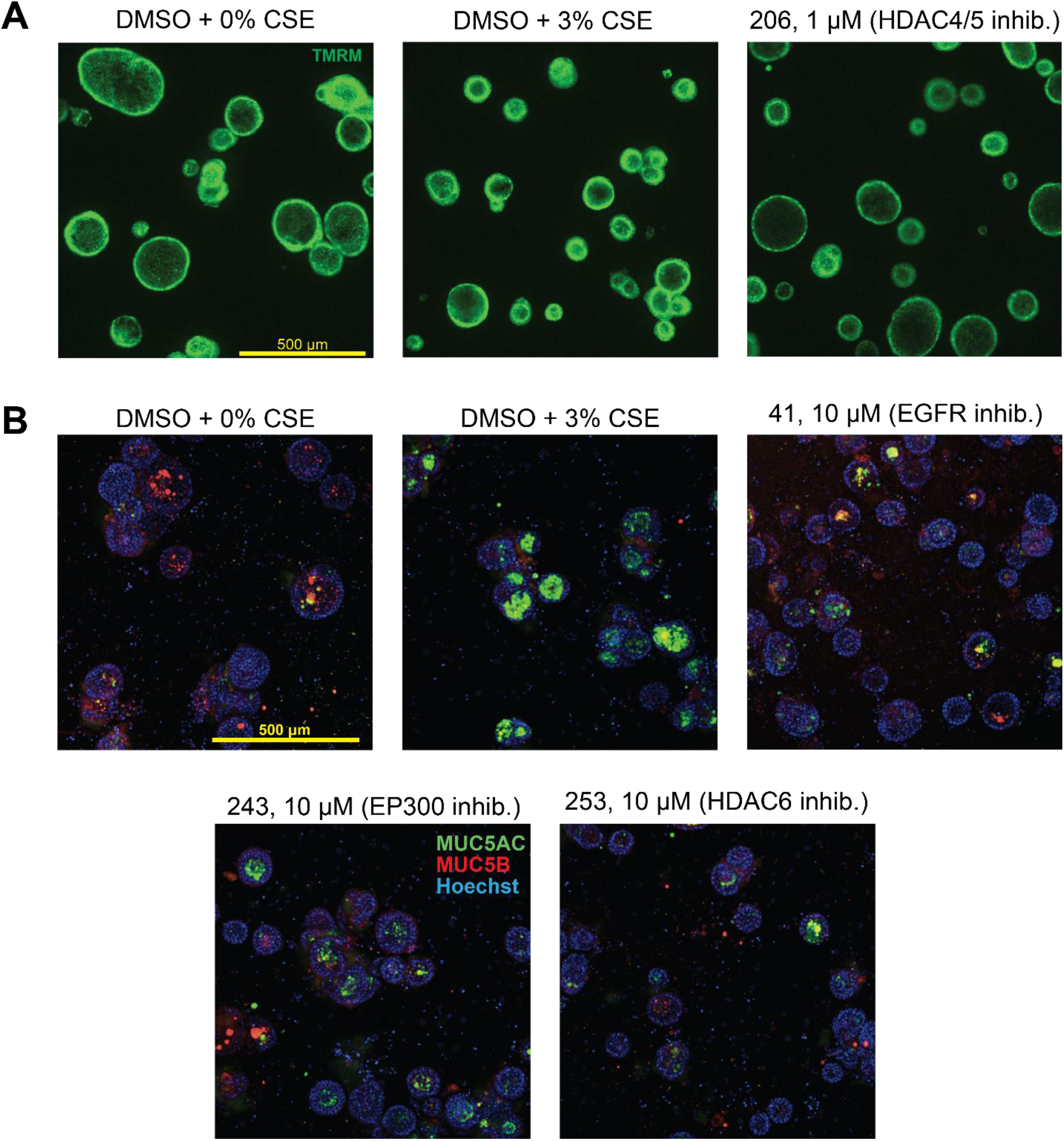
Attenuation of spheroid size decrease or MUC5AC increase by hit compounds. Final hit compounds (with hit concentration and corresponding target) that **A)** increased bronchosphere size compared to DMSO + 3% CSE control bronchospheres or **B)** decreased secreted MUC5AC. Mucus reduction hit compounds were further refined by manually comparing TMRM live dye-stained images of compound treated vs. control bronchospheres to observe aberrant phenotype that could indicate an unhealthy state. These images are shown in Supplemental Figure 5.

**Table 1:** Final list of hit compound from the validation screen and their corresponding targets.

| Hit Category | Compound # | Concentration (μM) | Public Identifier | Target |
| --- | --- | --- | --- | --- |
| **Swell** | 206 | 1 | LMK235(52) | HDAC4/5 |
| **Mucus** | 41 | 10 | AEE788(53) | EGFR |
| **Mucus** | 243 | 10 | CPI637(54) | CBP/EP300 |
| **Mucus** | 253 | 10 | CHEMBL3415627(55) | HDAC6 |

## Discussion

To discover effective therapies that target the underlying disease pathology for COPD, there is a pressing need for models that recapitulate the relevant disease biology with applicable readouts. Past attempts to create COPD models *in vitro* have not been able to capture both the complexity of cell differentiation prior to treatment or disease-relevant long term CSE exposure(12–15), or they lack the throughput necessary to effectively run drug discovery and target identification efforts(10,11). We have developed a high-throughput 384-well bronchosphere model which can mimic a COPD-like phenotype after long-term treatment with CSE. We were able to show that CSE-treatment for 1, 2, and 3 weeks results in a decrease in bronchosphere area and an increase in secreted MUC5AC in the bronchosphere lumen, both of which are phenotypes that have been previously shown in COPD-mimetic assay platforms and lend themselves to measurement by high-content images to provide quantitative high-throughput readouts of therapeutic efficacy for phenotypic screening(16,24).

While it is important to identify key phenotypes that are responsive to CSE exposure and suitable for high-throughput screening, it is equally important to ensure that we validate that the system mimics the changes that occur in humans due to long-term cigarette smoke exposure. We performed RNAseq of bronchospheres exposed to CSE for 1, 2, and 3 weeks and compared it against transcriptomic smoking signatures from three independent human datasets. We were able to show that genes upregulated in our bronchosphere model were enriched in the upregulated genes in all human smoking datasets, while genes downregulated in our system were also enriched in the downregulated genes across all smoking signatures from human samples. When looking at the top 75 upregulated and top 75 downregulated genes in the Tilley-1 comparison, we found that the direction of expression changes in our bronchosphere model was consistent with the expression changes in the human smoking datasets. Downregulated genes in our *in vitro* system were enriched in pathways and processes associated with EMT, ECM organization, and cell motility, suggesting that cell motility and cell-ECM interactions were negatively affected by CSE treatment. Of note, the downregulated genes were also enriched in pathways involved in cilium assembly and organization. Loss of cilia and downregulation in genes associated with ciliary cells were strongly associated with COPD(38,39), so this further indicates that the bronchosphere model recapitulates human disease states at the transcriptomics level. Upregulated genes were enriched in pathways involved in steroid(40) and xenobiotic metabolism(41), and these molecular functions were previously shown to be upregulated in smokers, suggesting increased metabolism upon CSE treatment. Upregulated genes were also enriched in pathways associated with oxidative phosphorylation and cellular response to oxidative stress, which were also associated with CSE treatment(41,42) and involved in the pathogenesis of COPD(43). The combination of phenotypic and transcriptomic changes occurring in the bronchospheres after CSE treatment provides a robust indication that the CSE-treated bronchospheres mimic the changes occurring in human smokers. Utilizing this model can accelerate drug discovery and benefit respiratory scientists with a translatable model system more consistent with human tissues in comparisons to animal models.

To demonstrate the screening potential of this system, we ran a small-molecule deck of 301 compounds of known, diverse mechanisms of action (MoA). Bronchospheres were treated with either DMSO or a compound of interest for 7 days, starting at day 7 of CSE exposure and ending at the 2-week endpoint. We were able to identify 26 hit compounds in the primary screen that either increased size or decreased MUC5AC in the lumen compared to control bronchospheres (DMSO + 3% CSE). Upon validation of the compounds, we were able to further reconfirm one hit that restored spheroid size (inhibitor of HDAC4/5) and three hits that reduced MUC5AC (inhibitors for EGFR, HDAC6, and CBP/EP300). Activation of EGFR(44) has been shown to contribute to COPD-associated phenotypes such as mucus overproduction and secretion, which would explain why inhibition reduces the MUC5AC levels within the bronchospheres lumen despite CSE treatment. Upregulation of HDAC6 has been implicated in ciliary shortening due to CSE treatment and has been shown to be upregulated in smokers both with and without COPD(45,46). However, the link between HDAC4, HDAC5, EP300 and COPD are not as established. Intriguingly, these proteins are involved in histone modifications and will be interesting to examine how CSE, as an environmental stimulus, may epigenetically modulate transcriptome and phenotypic changes using our bronchosphere model. Lastly, our initial pilot screen can be expanded to a larger follow-up screen, which may identify compounds with higher efficacy and specificity, resulting in better understanding of key drivers for CS-induced changed in COPD and better therapeutic solutions(47).

## Methods

### Tissue Culture

Normal human bronchial epithelial cells (HBECs) from 3 independent donors (Supplemental Table S1) were obtained from Lonza (CC-2540) and cultured in BEGM growth media (Lonza CC-3170), ranging in age from 12 to 38 years old. NIH/3T3 cells (ATCC CRL-1568) were cultured in 3T3 growth media consisting of DMEM/High Glucose (Hyclone SH30022.01), 10% FBS (Corning 35-015-CV), and 1% penicillin/streptomycin (HyClone SV30010). Bronchosphere differentiation media consists of 50% BEBM (CC-3171) and 50% DMEM/High Glucose supplemented with Bronchial Epithelial Cell Growth Medium SingleQuots™ (Lonza CC-4175).

To generate 3D bronchosphere cultures, 3T3s (100 cells/well) were seeded in 384-well PhenoPlates (Perkin Elmer CUSG02184) in 3T3 growth media and incubated overnight at 37°C, 5% CO_2_. 3T3 media was removed from the wells using a multichannel pipette and 10μL of 50% Matrigel (Corning 356230) in bronchosphere differentiation media was added to each well and incubated at 37°C for a minimum of 30 minutes to solidify the gel. HBECs were seeded at 100 cells/well in 30μL differentiation media with 5% Matrigel. The plates are incubated overnight at 37°C, 5% CO_2_ and 40μL of differentiation media per well was added the following day. Media changes occurred every 2-3 days; plate were aspirated down to 40μL per well and 40μL of fresh differentiation media was added on top. All trans-retinoic aid (Sigma R2625) was added fresh to differentiation media before each media change (50nM final concentration). Bronchospheres were cultured in differentiation media for a minimum of 14 days to encourage differentiation of distinct airway epithelial subtypes, as previously reported(24).

### CSE generation and treatment of bronchospheres

1R6F Small Batch cigarettes (Center for Tobacco Reference Products, University of Kentucky) were connected to a gas mixing chamber (Chemglass CG-1114-13) containing DMEM/high glucose. 100% CSE was generated by drawing smoke from 2 cigarettes per 10mL of DMEM at a continuous flow rate of 3.5 scfh, after which it was sterile filtered using a 0.22μm-pore vacuum filter.

### Live imaging and analysis of spheroid size

The day before the indicated timepoint, 5μL of 425nM TMRM dye (ThermoFisher, Cat# T668) in differentiation media was added to each well (final well concentration was 25nM) and the plates were incubated overnight. Images were acquired at 4X magnification using the ImageXpress confocal microscope (Molecular Devices).

Image segmentation is done using the open-source deep-learning based segmentation tool Cellpose (https://github.com/mouseland/cellpose). For spheroids segmentation, the Cellpose pretrained model “CP” is used with proper object diameter parameter, which is roughly equals to average spheroid sizes of a specific experiment run. Segmentation results are manually inspected to ensure reasonable segmentation quality and the diameter value was adjusted when necessary, in rare cases. Ilastik (https://www.ilastik.org/), the open-source interactive learning toolkit is then used to calculate features for each spheroid such as size and total signal intensity, using both the raw image and segmentation results from Cellpose as import, an object classification model with one class classification was built and applied to the entire dataset plate-by-plate using the batch mode. Spheroids features calculated from each image are then assembled into TIBCO Spotfire (https://www.tibco.com) and final views showing well-level aggregated values such as spheroids size or intensity were plotted and calculated within Spotfire for further analysis. Spheroid areas were normalized to the average CSE 0% value per donor for each time point, to account for donor-to-donor size differences.

### Fluorescent imaging and analysis of luminal mucus content

On timepoint day, plates were aspirated down to 30μL and 30μL of 8% paraformaldehyde (Electron Microscopy Sciences 15714-S) in PBS was added per well. Plates were fixed for 1 hour then washed three times with PBS. Wells were blocked with blocking buffer—final concentration 0.1% bovine serum albumin (Milipore Sigma A7030), 0.2% Triton-X (Fisher Scientific BP151-100), 0.04% Tween-20 (Fisher Scientific BP337-100), 10% goat serum (Gibco 16210) in PBS—and incubated for 1.5 hours. Plates were then aspirated to 40μL and 10μL of primary antibodies diluted 1:100 in blocking buffer was added to each well (final dilution 1:500) and incubated overnight at room temperature. Primary antibodies used were: MUC5AC (Thermo MA5-12178), MUC5B (Sigma HPA008246-100UL), α-acetylated tubulin (Sigma T6793-.2ML). Plates were washed three times with PBS with 0.2% Triton-X and 0.04% Tween-20 (IF wash buffer), then 10μL of secondary antibodies diluted 1:100 and Hoechst (Invitrogen H3570) diluted 1:1000 (final dilutions 1:500 and 1:5000) was added to each well and incubated for 4 hours at room temperature. Secondary antibodies used were: AlexaFluor 488 goat anti-mouse (Invitrogen A11001) and AlexaFluor 647 goat anti-rabbit (Invitrogen A21245). Plates were washed three times with IF wash buffer then three times with PBS. Plates were imaged at 5X magnification using the Opera Phenix High Content Screening System (Perkin Elmer). 2D maximum projection(MIP) images are generated from raw z-stack images to streamline downstream analysis.

Segmentation is done using the open-source deep-learning based segmentation tool Cellpose (https://github.com/mouseland/cellpose). For spheroids segmentation, the Cellpose pretrained model “CP” is used with proper object diameter parameter, which is roughly equals to average spheroid sizes of a specific experiment run. Segmentation results are manually inspected to ensure reasonable segmentation quality and the diameter value was adjusted when necessary in rare cases. Ilastik (https://www.ilastik.org/), the open-source interactive learning toolkit is then used to calculate features for each spheroid such as size and total signal intensity, using both the raw image and segmentation results from Cellpose as import, an object classification model with one class classification was built and applied to the entire dataset plate-by-plate using the batch mode. In addition to the original spheroid channel, two more mucus antibody-specific staining channels are also imaged. The total mucus-specific intensity and the area occupied within each spheroid from each additional image channel need to be calculated. To do that we rely on CellProfiler (https://cellprofiler.org/), an open-source cell image analysis software. A CellProfiler pipeline was built by segmenting mucus active area from both mucus channels within each spheroid mask generated by Cellpose in previous step, then the percentage of mucus active area and the total intensity in each spheroid are calculated. Spheroids features calculated from both Ilastik object classification and CellProfiler pipeline for each image are then combined and assembled into TIBCO Spotfire (https://www.tibco.com) and final views showing well-level aggregated values of mucus percentage occupation and total signal inside spheroids were calculated and plotted within Spotfire for further analysis.

### RNA isolation and bulk RNA sequencing

At all timepoints, RNA was isolated from 0 or 3% CSE-treated bronchospheres using the Qiagen RNeasy Mini Kit (Qiagen 74106). The RNA yield was measured using Agilent TapeStation (4200 TapeStation System) at 260 nm. 150 ng of total RNA was used to make RNA sequencing libraries using the TruSeq Stranded mRNA Prep Kit (Illumina 20020595). Libraries were run on a 50-cycle NovaSeq (SP flowcell, 2 lanes) sequencing run and yielded an average sequencing depth of 15 million reads per sample. Reads were generated from a NovaSeq instrument with bcl2fastq from Illumina, v 2.20.0.422-2 and were aligned to a combined human/mouse genome provided by 10xGenomics, version 2020-A with STAR aligner v2.7.3a(48). Gene counts were assessed with RSEM v1.3(49). and were normalized using DESeq2 v.1.28.1, in R version 4.0(50). The output was log2(x+1) normalized.

### Smoking signatures

GEO datasets comparing smoking effects in human cohorts were examined and data from GSE20257, GSE7895, and GSE63127 were acquired. GSE20257 had GPL570 array data for 53 healthy non-smokers, 59 healthy smokers, and 23 COPD smokers(25). GSE7895 had GPL96 array data for 21 never smokers, 31 former smokers, and 52 current smokers(26). GSE63127 had GPL570 array data for 87 non-smokers and 143 smokers(27). Log2 transformed data were examined and linear regression was performed to identify smoking signatures using different comparison groups. The models controlled for age and sex.

### Primary compound screen and hit analysis

A detailed schematic of the screen set-up is shown in Figure 3. Bronchospheres were differentiated for two weeks, after which they were treated with either 0 or 3% CSE for an additional 2 weeks. On day 7 of CSE treatment, bronchospheres were treated with either DMSO or compounds from the MoA deck at 1 or 10μM concentrations. On day 11, the bronchospheres were re-treated with compounds and cultured until day 14 (2 weeks) of CSE treatment. On day 13, 5μL of 425nM TMRM dye (ThermoFisher, Cat# T668) in differentiation media was added to each well (final well concentration was 25nM) and the plates were incubated overnight. Images were acquired at the 2-week endpoint (day 14 of CSE treatment) at 4X magnification using the ImageXpress confocal microscope (Molecular Devices) to observe changes to spheroid size due to compound treatment. Bronchosphere plates were then fixed with paraformaldehyde and stained for MUC5AC, MUC5B, and nuclei to observe changes in luminal MUC5AC due to compound treatment (see previous section *“Fluorescent imaging and analysis of luminal mucus content*”).

For swell (spheroid size increase) hits, average spheroid size per well was normalized to the median spheroid area of DMSO + 3% CSE treated bronchospheres for each plate. Compounds needed to have a fold change that was greater than 2 * (Median Absolute Deviation [MAD] of DMSO + 3% CSE condition) to be considered hits. To filter out compounds that appeared to cause toxicity, number of spheroids per well needed to be within 3 * MAD of DMSO + 0% CSE, which had lower counts than the 3% CSE and was therefore used as the healthy cut off. Similarly for mucus reduction hits, average MUC5AC ratio per well was normalized to the median MUC5AC ratio of DMSO + 3% CSE treated bronchospheres for each plate and converted to a Log2(Fold Change). Compounds needed to have a −Log2FC that was greater than 2 * (MAD of DMSO + 3% CSE condition) to be considered hits. To filter out compounds that appeared to cause toxicity, total nuclear intensity per well needed to be within 3 * MAD of DMSO + 3% CSE, which had lower intensity than 0% CSE and was therefore used as the healthy cut off. Additionally, the spheroid counts per well from the swell data was further used to determine toxicity in the mcus hits. A compound was considered a *swell hit* if at least 2 well replicates had a fold change greater than 2 * MAD of DMSO + 3% CSE, all three well replicates had spheroid counts within 3 * MAD of DMSO + 0% CSE, and if at least two of the wells did not increase MUC5AC ratio by greater than 2 * MAD above DMSO + 3% CSE. A compound was considered a *mucus hit* if at least 2 well replicates had both a −Log2FC greater than 2 * MAD of DMSO + 3% CSE and a total nuclear intensity within 3 * MAD of, all three well replicates had spheroid counts within 3 * MAD of DMSO + 0% CSE, and if it did not decrease bronchospheres area by greater than 2 * MAD below DMSO + 3% CSE. The results of the primary screen are showing in Figure 3 and the data associated with the scatter plots is in Supplemental Table S8. The box plots for the DMSO treated bronchospheres that were used as cutoffs for the screen are shown in Supplemental Figure 6. While 28 compounds were identified as hits, two of the compounds (Compound# 144 and 196, refer to Supplemental Table S7 for compound information by numerical code) were not internally available in a second batch for validation and were therefore excluded from the validation screen, resulting in 26 hit compounds.

### Secondary hit validation screen and analysis

The 26 compounds that were hits from the primary screen were tested in a secondary validation screen with the same experimental set-up as the primary screen but different batches of the compounds (not the exact batch as was used in the primary screen). Compounds were tested at 1 and 10μM concentrations, with compound treatment at day 7 and 11 of CSE treatment, with the assay endpoint at day 14 (2 weeks) of CSE treatment. Spheroids were imaged live with TMRM for swell/spheroid size changes, then fixed and stained for MUC5AC ratio changes. Average spheroid area and MUC5AC ratio raw values were plotted in GraphPad Prism and a one-way ANOVA with Dunnett’s multiple comparisons test was run to identify validated hits.

### Statistical analysis for RNA sequencing analysis

Because human bronchial epithelial cells in the experiment were based on three subjects (Supplemental Table S1), principal component analysis (PCA) was performed to assess donor effects. First and second principal components (PCs) were capturing donor effects such as alcohol status, age, and sex. Cigarette smoking excrement (CSE) related signals were captured in the third and fourth PCs. Therefore, mixed model utilized to identify differential expressed genes (DEGs) accounting for donor status as random effects. After testing the model sex effect was fully accounted by Donor variable and the following model was implemented for each time point: *Gene expression* ~ 1 + *alcohol status* + *Age* + (1\*Donor*). PCA was performed again to verify that the first PC, the major source of variation, is from CSE and second largest variation is due to weeks of CSE treatment (Supplemental Figure 2). To maximize power to identify CSE dependent DEGs, analysis pooling all time points were conducted with the following model: *Gene expression ~ 1* + *days treated* + *alcohol status* + *Age* + (1|*Donor*). Benjamini-Hochberg procedure was performed to correct for multiple testing. Gene-Set Enrichment Analysis (GSEA) was performed using fgsea package in R(51). Minimum and maximum gene-set size were 5 and 500, respectively, and statistical inference was made with 50,000 permutations. All association results were corrected for multiple-testing using Benjamini-Hochberg procedure and significant associations were controlled at 5% FDR. Significantly associated pathways with gene-set overlap size larger than 30 genes were selected for visualization.

### Statistical analysis for all other graphs

Unless otherwise indicated, statistical analysis was performed using GraphPad Prism9, and the threshold for significance was set at p < 0.05. Detailed information on the statistical analysis of each figure is written in the corresponding figure legend.

## Acknowledgements

The authors would like to thank Dr. Fred King, Dr. Barun Okram, Dr. Paul Rucker, Dr. Dean Phillips, Dr. Jimmy Elliot, Dr. Kevin White, Khaushik Subramanian, Stephen Attle, Dr. Michael O’Sullivan, and Dr. David Rowlands for helpful discussion and/or guidance in development of methodology or experimental design.

## Author Declarations

### Author contributions statement

Conceptualization: P.B., W.B., E.T.; Methodology: P.B., K.S., D.K., D.Q.; Data Generation and Analysis: P.B., Y.J.W., K.S., K.C., S.W.B., O.R.; Writing Manuscript: P.B., Y.J.W., K.C.; Supervision: B.F., J.W., W.B., E.T. Funding: Novartis Institute of Biomedical Research. All authors reviewed and approved this manuscript.

### Data availability

The RNAseq data reported in this paper will be uploaded (and the accession number provided) prior to paper acceptance. The data that supports the results in the paper are available in the article or supplementary information. Additional data for the figures are available from the corresponding author upon reasonable request.

---

**Supplemental Figure 1:**
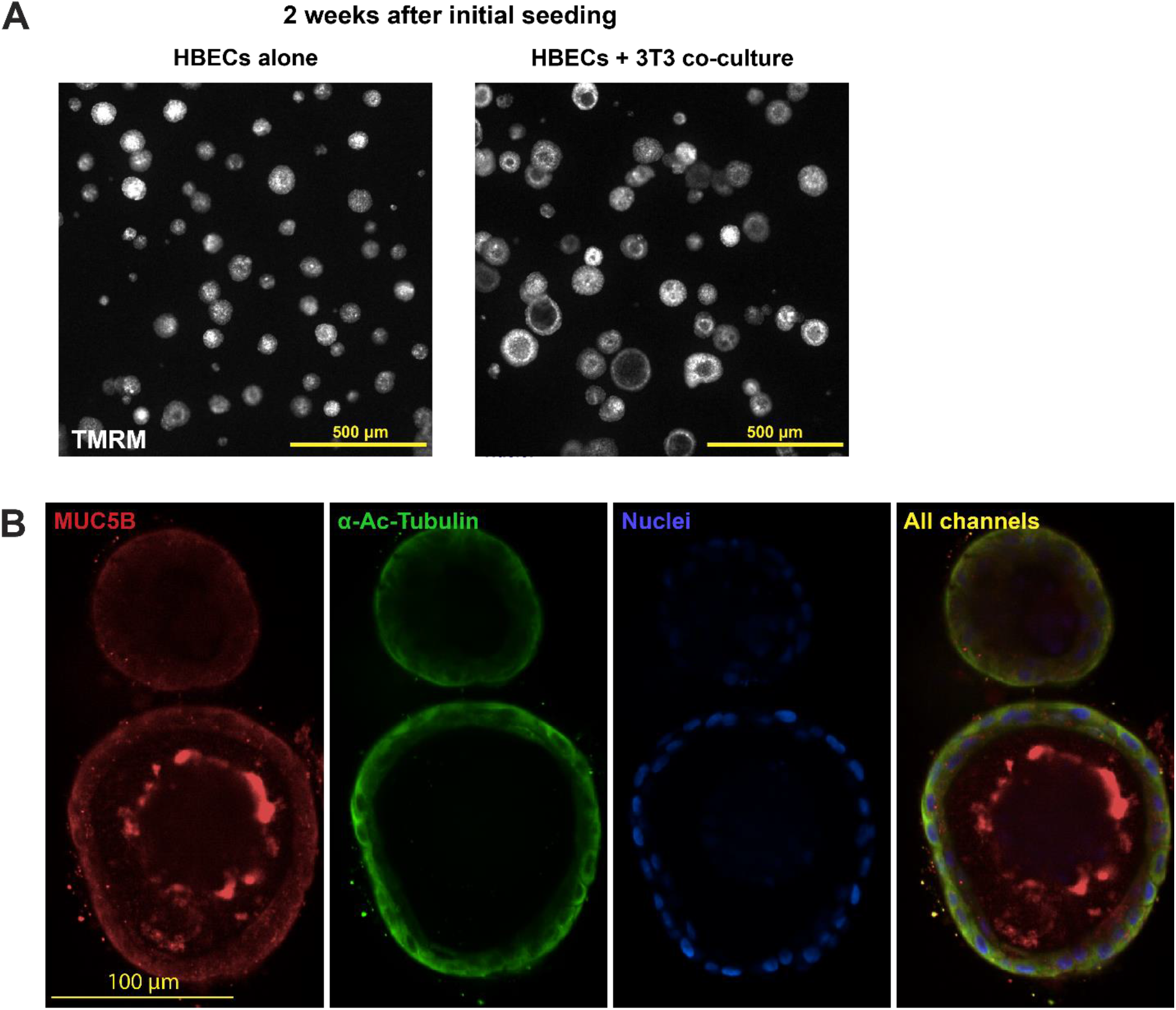
Bronchosphere differentiation. **A)** Bronchospheres 2 weeks after initial seeding, with and without 3T3 feeder layer. **B)** After two weeks, bronchospheres stain positive for MUC5B and α-acetylated tubulin, indicating that they have differentiated to contain secretory and ciliated cell subtypes, as previously shown(18,24).

**Supplemental Figure 2:**
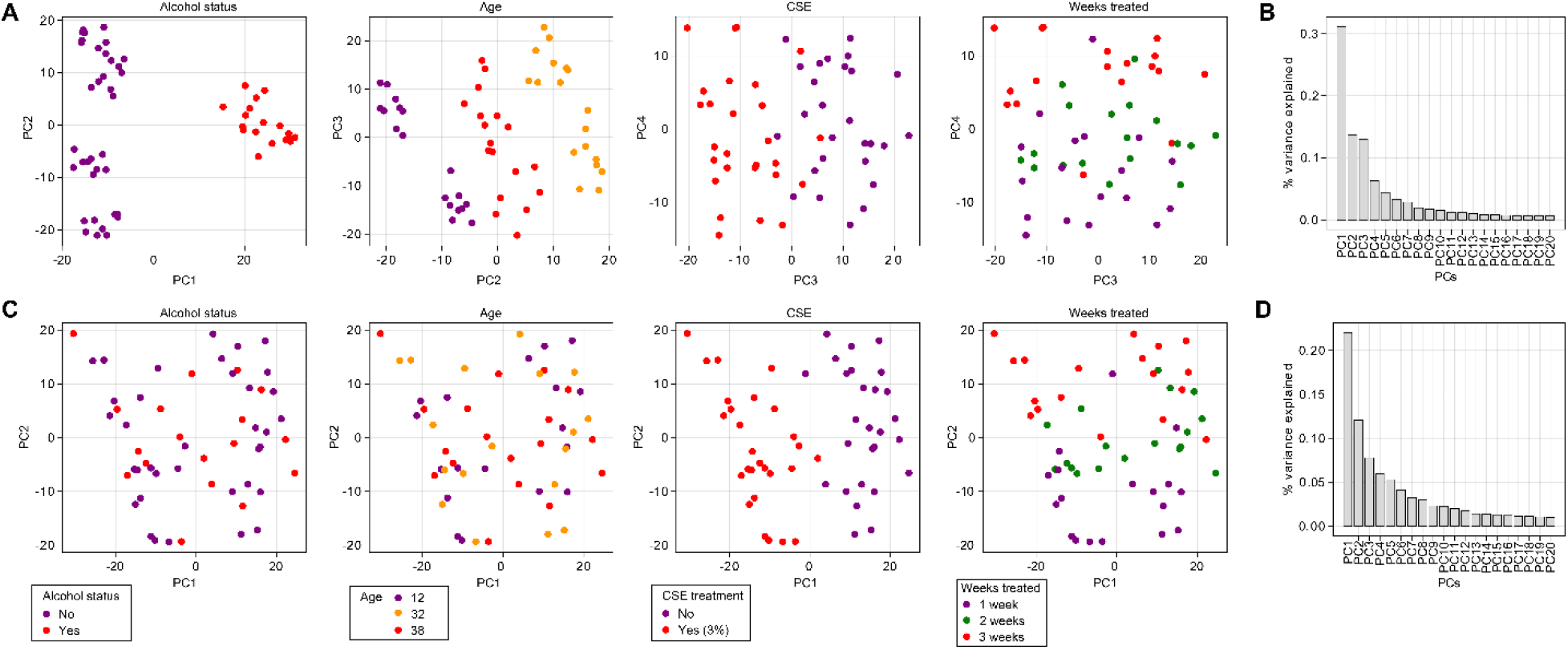
PC effects after DESeq2 only normalization vs mixed model. **A)** Scatter plots visualizing PC effects after DESeq2 normalization only. **B)** Bar plot describing the % of variance explained by each PC after DESeq2 normalization only. The major PCs capture donor specific effects, meaning that the most variance in the data is not reflective of CSE treatment effect. **C)** Scatter plots visualizing PC effects after removing the donor, age, and alcohol status using mixed model. The major PCs capture CSE treatment effect. **D)** Bar plot describing the % of variance explained by each PC after covariate adjustment using mixed model.

**Supplemental Figure 3:**
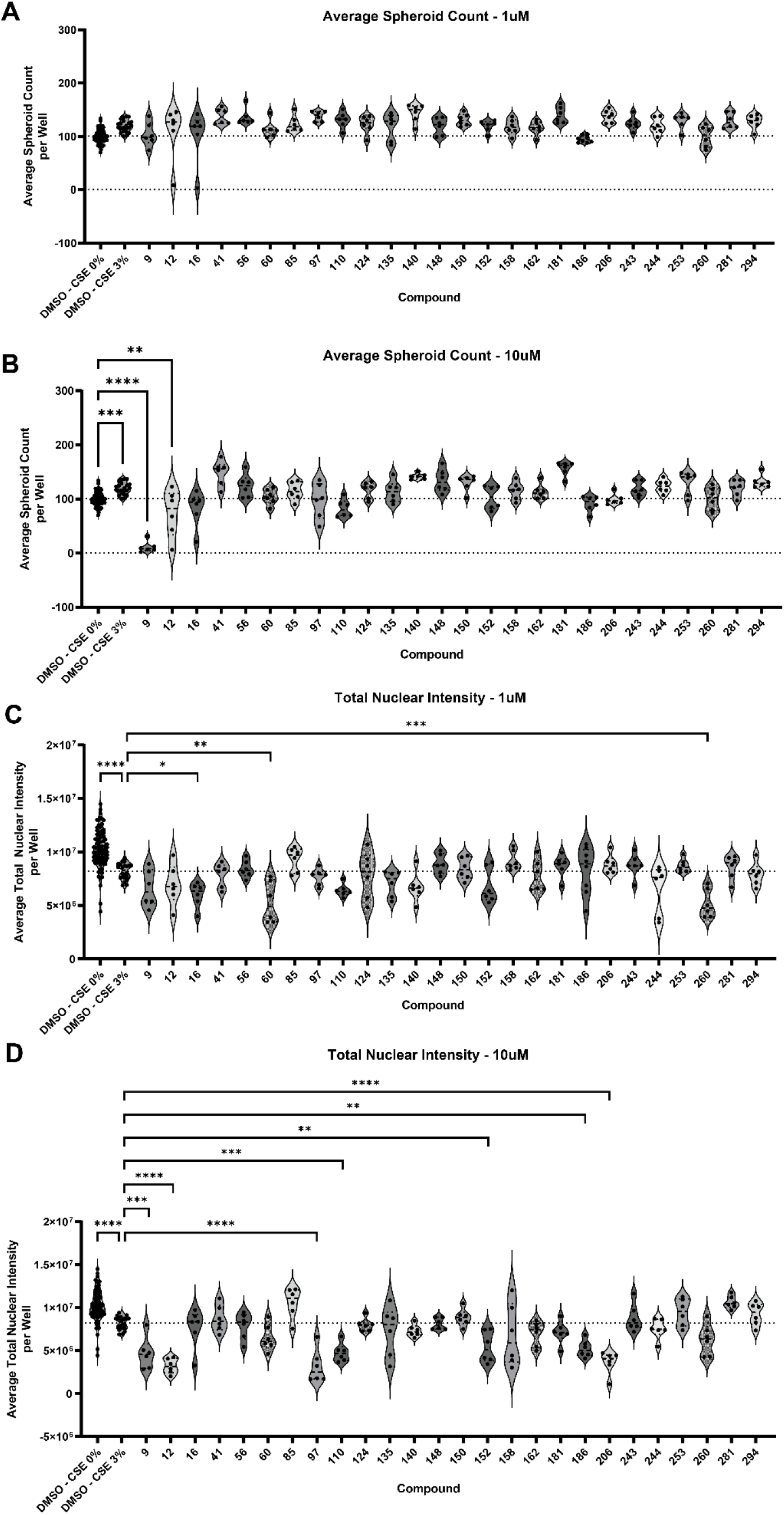
Utilizing reduction in spheroid counts and/or total nuclear intensity to determine toxic effects of compounds. Compounds that were identified in the primary screen as modulators of both phenotypes were tested at 1μM and 10μM. The number of spheroids per well (spheroid counts) was measured from the spheroid size/swell readout with TMRM live dye staining, while the total nuclear intensity per well (total nuclear intensity) was measured from the MUC5AC ratio readout with Hoechst staining after fixation. Spheroid counts per well were calculated for compound treatment at **A)** 1μM and **B)** 10μM concentrations. Total nuclear intensity per well was calculated for compound treatment at **C)** 1μM and **D)** 10μM concentrations. Compounds that were found to significantly decrease counts or nuclear intensity were considered toxic and filtered out of final hit selection. They are indicated appropriately in Figure 4. All individual data points represent biological replicates. All plots were analyzed by ordinary one-way ANOVA with Dunnett’s multiple comparisons test. *p<0.05; **p<0.01, **p<0.01, ***p<0.001, ****p<0.0001.

**Supplemental Figure 4:**
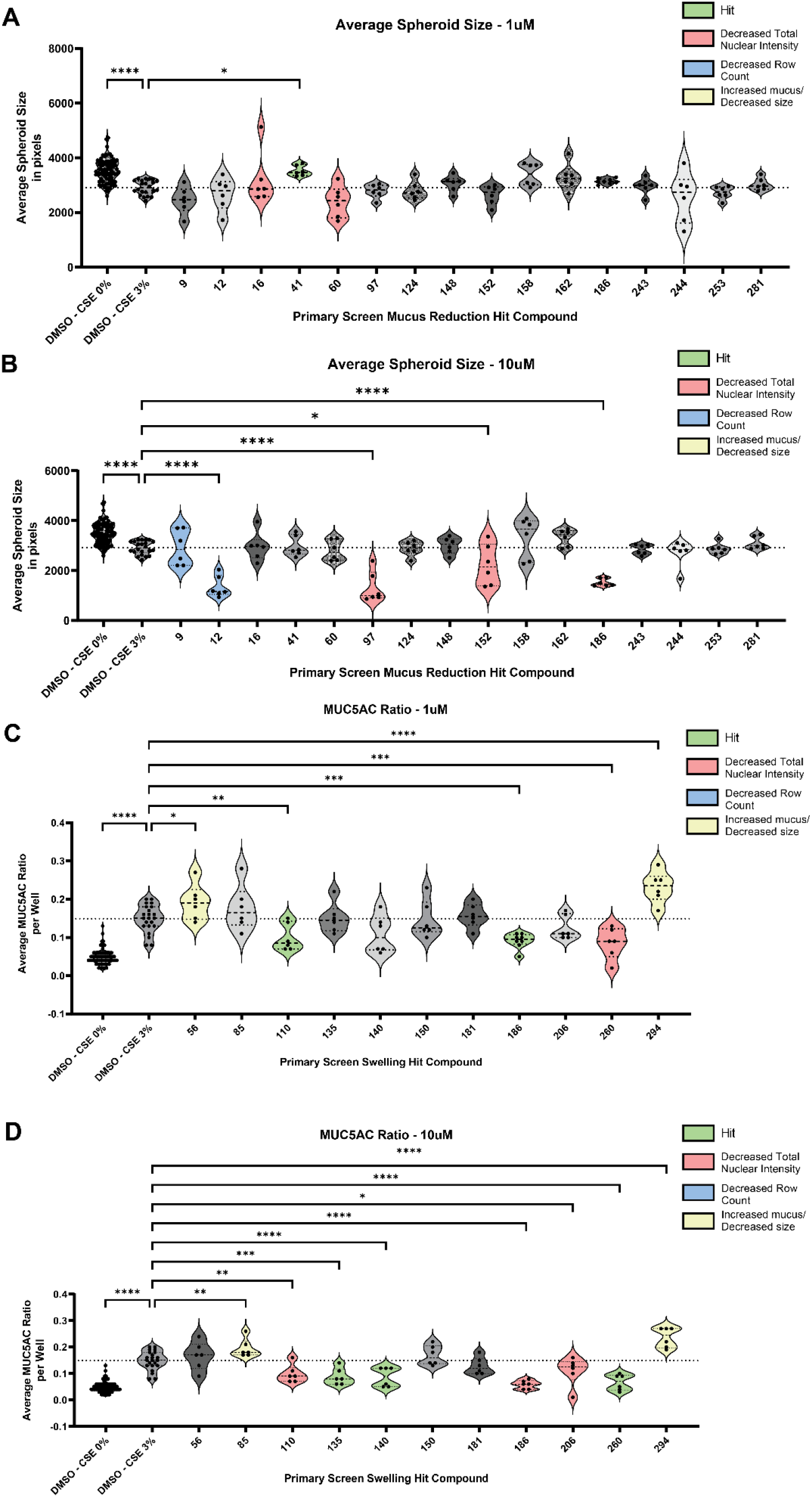
Spheroid size changes due to primary mucus reduction hits and mucus changes due to primary swell hits. Compounds that were identified in the primary screen as modulators of *MUC5AC reduction* were tested at **A)** 1μM and **B)** 10μM concentrations to see if they affect <u>spheroid size</u>. No compounds caused significant shrinkage that were not already identified through the toxicity readouts in Supplemental Figure 3. Compounds that were identified in the primary screen as modulators of *spheroid size/swell* were tested at **C)** 1μM and **D)** 10μM concentrations to see if they affect <u>MUC5AC ratio</u>. Compound 186 (1μM) is the only compound that was a primary hit in both swell and MUC5AC ratio and is plotted in both readouts in both Figure 4 and Supplemental Figure 4 and is identified as a hit compound in Figure 4C. Compounds 56 (1μM), 85 (10μM), and 294 (1 and 10μM) significantly increased MUC5AC ratio compared to DMSO + CSE 3% control and were therefore filtered out of swell hit selection. These compounds are appropriately labeled in Figure 4. Hits (colored green) identified in **A)-D)** were not selected as final hits since they did not produce the desired effect in the primary screen (Figure 3). All individual data points represent biological replicates. All plots were analyzed by ordinary one-way ANOVA with Dunnett’s multiple comparisons test. *p<0.05; **p<0.01, **p<0.01, ***p<0.001, ****p<0.0001.

**Supplemental Figure 5:**
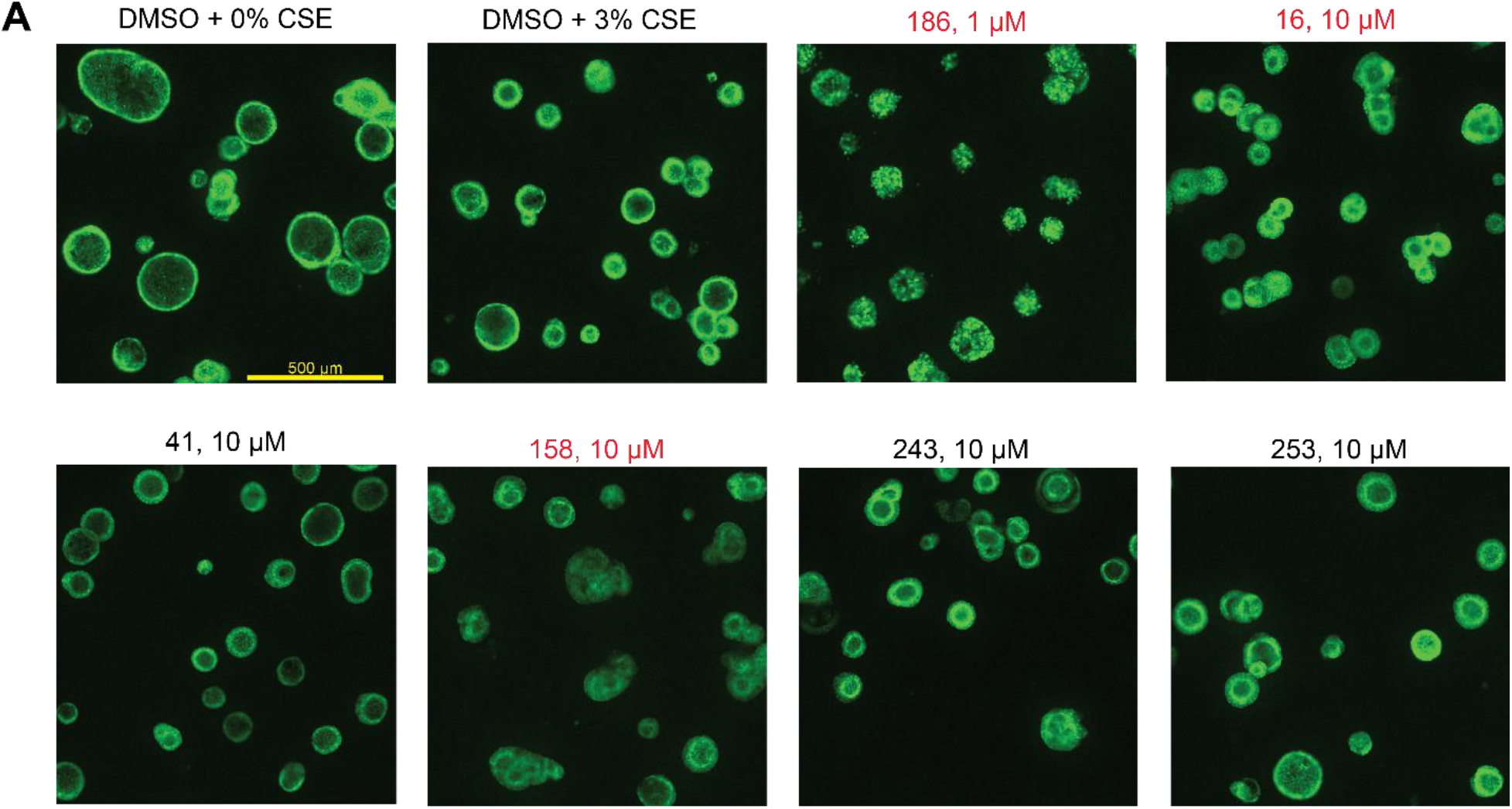
Refinement of mucus reduction hits via visual confirmation of live bronchosphere images. **A)** TMRM live dye-stained images of bronchospheres treated with hit compounds and concentrations. The images reveal that Compound 186 (1μM), 16 (10μM), and 158 (10μM) cause phenotypic changes to the bronchospheres that indicate an unhealthy state compared to DMSO + 3% CSE control, such as rough spheroid boundaries or loss of a clear lumen (indicated in red). These compounds have thus been filtered out of the final hit list.

**Supplemental Figure 6:**
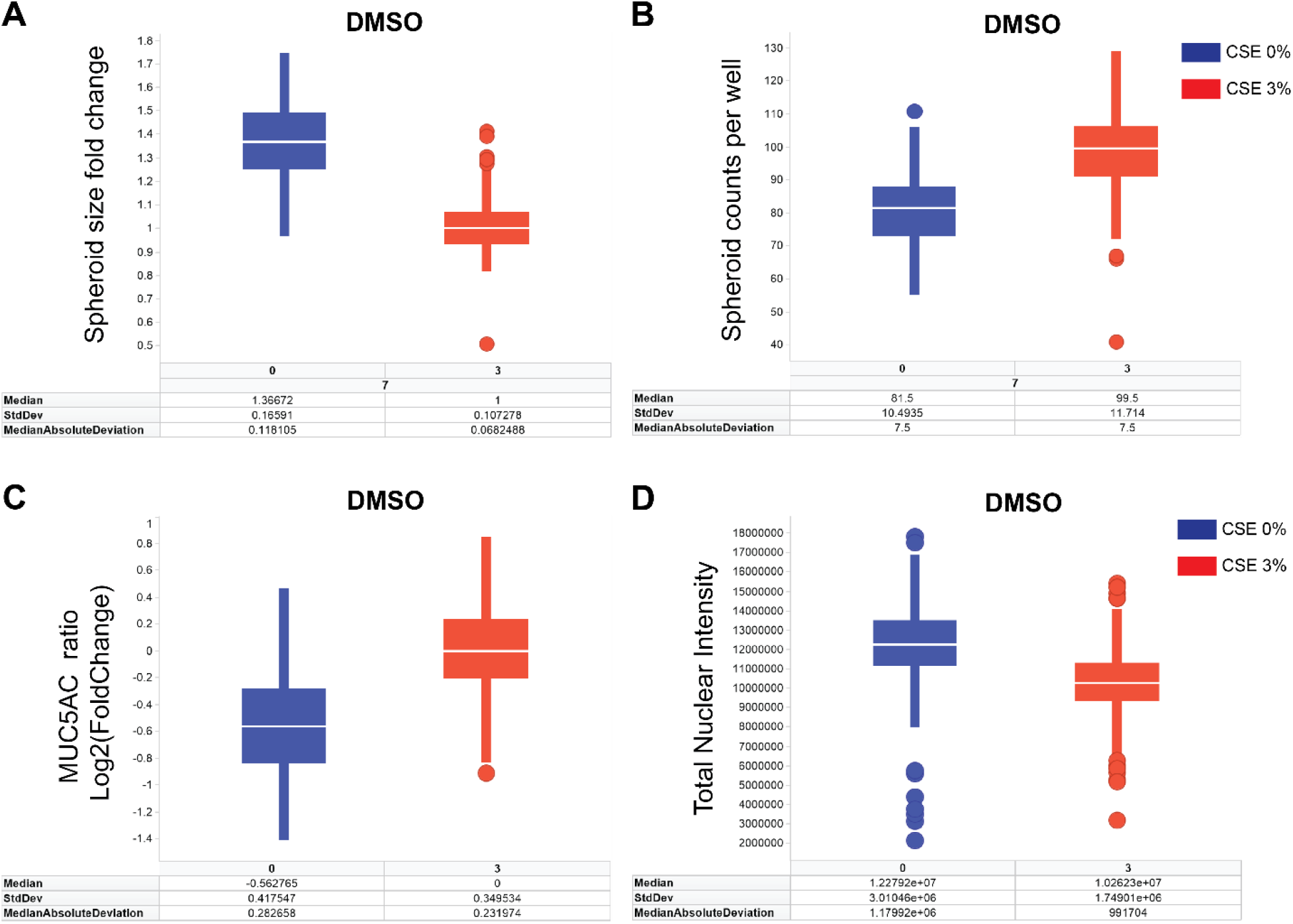
Median absolute deviation values of DMSO-treated bronchospheres (CSE 0 and 3%) used for hit-picking cutoffs. DMSO-treated spheroids at both 0 and 3% CSE were plotted after plate normalization (see details in *“Primary compound screen and hit analysis”* section of Methods) to obtain the median absolute deviation (MAD) for **A)** the spheroid size fold change and **B)** the spheroid counts per well from the spheroid size assay, and **C)** the MUC5AC ratio Log2(Fold Change) and **D)** the total nuclear intensity from the MUC5AC reduction assay.

